# Transcriptional drift in aging cells: A global de-controller

**DOI:** 10.1101/2023.11.21.568122

**Authors:** Tyler Matsuzaki, Corey Weistuch, Adam de Graff, Ken A Dill, Gábor Balázsi

## Abstract

As cells age, they undergo a remarkable global change: In *transcriptional drift*, hundreds of genes become overexpressed while hundreds of others become underexpressed. Using archetype modeling and Gene Ontology analysis on data from aging *Caenorhabditis elegans* worms, we find that the upregulated genes code for sensory proteins upstream of stress responses and downregulated genes are growth- and metabolism-related. We propose a simple mechanistic model for how such global coordination of multi-protein expression levels may be achieved by the binding of a single ligand that concentrates with age. A key implication is that a cell’s own responses are part of its aging process, so unlike for wear-and-tear processes, intervention might be able to modulate these effects.

## Introduction

Upon aging, cells can undergo changes that are either *extrinsic* to the cell (non-autonomous), including signaling between tissues or *intrinsic* to the cell (autonomous). Cell-intrinsic factors can be roughly classified into two types: either (1) *wear-and-tear*, or (2) *the cell’s responses*, i.e., adaptive actions taken by the cell in response to aging. Examples of wear-and-tear include when mitochondria become less effective [1, 2], membranes become leaky [3, 4, 5], DNA, lipids, and proteins accumulate damage [6, 7, 8], and protection within the proteostasis system weakens [9, 10, 11].

A manifestation of aging is changes in gene expression. On the one hand, with some notable exceptions [12], aging can be associated with increases in transcriptional noise, which is the cell-to-cell variation in gene expression and which results in variations in mRNA and protein levels [13, 14, 15].

On the other hand, of interest here, Rangaraju *et al*. have recently explored more systematic changes in gene expression in aging *Caenorhabditis elegans* worms, which they call *transcriptional drift*. In transcriptional drift, hundreds of genes become increasingly overexpressed with age (relative to younger cells) while hundreds of others become increasingly underexpressed within the same cell [14]. Transcriptional drift has been observed not only in *C. elegans*, but also within mice and humans [14, 16]. Transcriptional drift is interesting because it may arise primarily as an actionable cell response to aging and thus potentially be susceptible to intervention. In support of this notion, it has been found that inhibiting transcriptional drift extends the worm’s lifespan [14]. And, while similar large-scale concerted changes in gene expression occur in the Environmental Stress Response (ESR) [17] in yeast and in the Integrated Stress Response (ISR) [18] in worms and higher organisms to combat stress, the ESR and ISR are typically only transient, whereas transcriptional drift is prolonged and persists over the full process of aging.

In the present work, we take three steps to analyze the transcriptional profile data of *C. elegans* over time, from Rangaraju *et al*. [14]. First, using Normalized Nonnegative Matrix Factorization (N-NMF) to analyze patterns in the data in an unbiased way, we identify two underlying archetypes that capture this concerted transcriptional variation with age. Second, we use gene ontology (GO) analysis to determine which cell functions are involved in these archetypes, i.e. which functions are up- and down-regulated in aging. Third, we propose a simple biophysical model to explain how such many-protein coordination could be achieved in a simple way.

## Gene grouping by archetype analysis

To identify concerted temporal signatures of gene expression in the data, we applied Normalized Nonnegative Matrix Factorization to the data of Rangaraju *et al*. [14]. NMF is a widely-used clustering algorithm for decomposing high-dimensional nonnegative signals into their dominant constituent parts [19]. Somewhat like Principal Component Analysis, these component parts or *archetypes* represent coupled collections of signals that, roughly, behave the same way. However, this approach has two key advantages. First, the components tend to cluster the signals into distinct parts [19, 20]. Second, by adding a normalization constraint, our NMF approach gives the relative contributions of the parts to each data sample. This method, which we detail in the SI, has recently been used to identify and to score the enrichment of distinct functional modules in many-gene cancer expression data [21, 22]. Our treatment allows us to find patterns within the transcriptional data in an unbiased manner and evaluate how the components of these patterns evolve over time.

One principal finding is that the aging *C. elegans* data is best represented by two dominant *archetypes* (see Figure A1 in the SI). Each archetype is a grouping of hundreds of genes. An archetype can be thought of as an idealized exemplar, a kind of functional averaging over types of proteins, that best characterize the behavior (increasing or decreasing with age) within the group. In our data set, the archetypes that emerged were genes that had either monotonically increasing or monotonically decreasing expression levels measured in counts per million (cpm) over time. The relative contributions of these two archetypes to the total *C. elegans* gene expression varies over time (see Fig 1) and represents the concerted transcriptional changes associated with aging.

**Figure 1.**
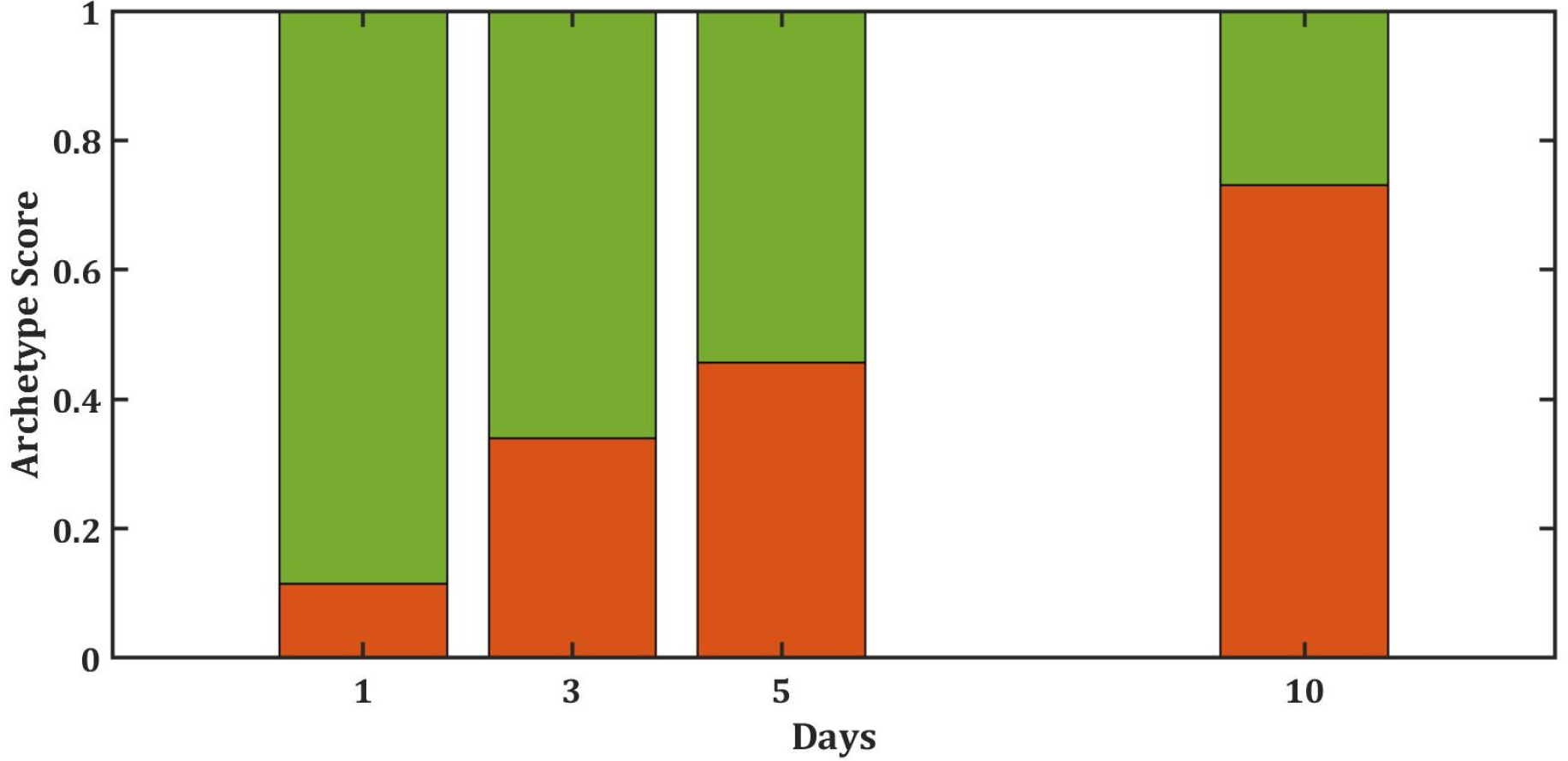
The relative changes of the two archetypes of genes with age. Using normalized nonnegative matrix factorization, we identified two key archetypes: one that increases with age (red) and one that decreases (green).

To gain insight into the sources of these archetypes, we extracted and examined the concerted time-dependent behaviors of the most representative genes in the relative composition of the *C. elegans* archetypes. Figure 2A shows a histogram of correlations among gene expression levels in *C. elegans*. The figure shows the numbers of genes for which expression tends to go down (left) or up (right), as a function of age. A remarkably large fraction of the whole genome changes systematically with age – either up or down – as seen by the areas under the curve of the two peaks on the left and right. We focus on these two sets, since they dominate the changes in proteome composition. We defined these dominant components as having a Pearson correlation coefficient with the global archetypes of *≤* −0.9 or *≥* 0.9, for a total of 1859 downregulated and 3006 upregulated genes, respectively. These dominant components are indicated to the left and right of the purple lines in Figure 2A. The expression of these genes are plotted in 2B. Note that the upregulated genes tend to increase roughly linearly with age, while the downregulated ones tend to decrease following a saturating function, like a Michaelis-Menten binding process.

**Figure 2.**
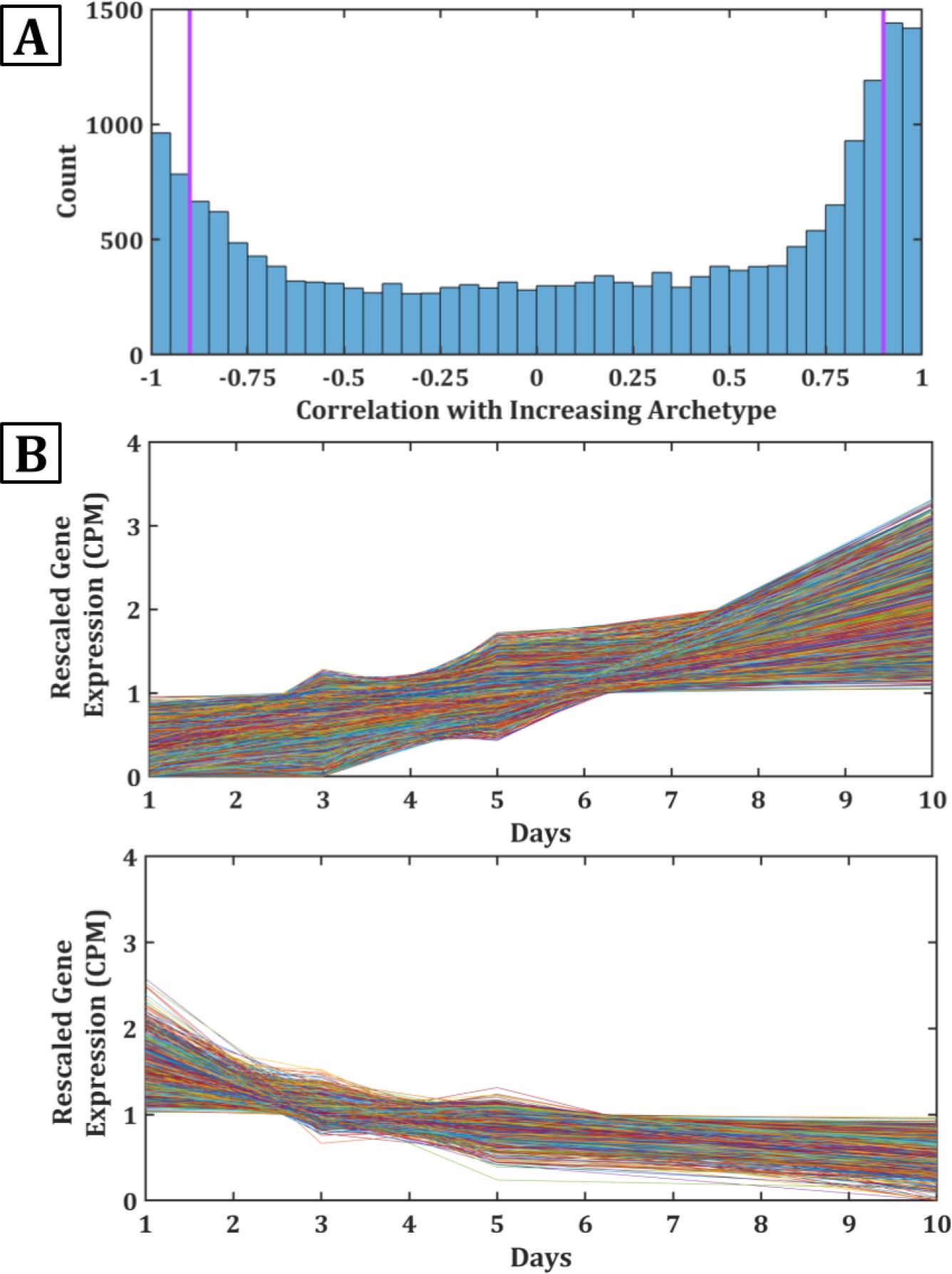
Much of the transcriptome changes with age. (A) Pearson correlation coefficient of all genes vs the two archetypes.Genes with high positive *R*^2^ are strongly monotonically increasing whereas genes with high negative *R*^2^ are monotonically decreasing. Genes selected as archetype centers are to the left and right of the purple lines. (b) Rescaled expression data for genes with correlation coefficients *≥* 0.9 (top) and *≤* −0.9 (bottom).

## Gene Ontology analysis of protein functions

The observed transcriptional drift appears to be non-random; there is a patterning in terms of which protein functionalities go up and which go down. To determine what biological functions are systematically upregulated and downregulated with age, we analyzed each subset using PANTHER Gene Ontology (GO) Enrichment Analysis for functional classifications with Fisher’s Exact Test [23, 24, 25]. This allowed us to compare which genes were overrepresented in each subset compared to our reference list (consisting of all genes in the full data set). Table 1 and SI Data give details of the GO analysis.

**Table 1:** Examples of GO categories in archetypes.

| Gene Ontology | Fold Enrichment | P Value | Change |
| --- | --- | --- | --- |
| Chemical Synaptic<br>Transmission, Postsynaptic | 2.52 | 7.60E-06 | Upregulated |
| Nervous System Processes | 2.01 | 1.87E-12 | Upregulated |
| G Protein-Coupled<br>Receptor Activity | 1.58 | 1.16E-12 | Upregulated |
| Cell Communication | 1.24 | 1.00E-7 | Upregulated |
| Proton Motive Force-Driven<br>ATP Synthesis | 4.09 | 2.32E-3 | Downregulated |
| TCA Cycle | 3.45 | 3.24E-03 | Downregulated |
| mRNA Processing | 2.93 | 8.34E-11 | Downregulated |
| Protein Ubiquination | 1.85 | 2.04E-6 | Downregulated |

We found that the upregulated archetype is enriched in functions related to sensing and transmitting signals. Genes having increased expression include acy-2, an adenylyl cyclase, and str-88, a GPCR protein and genes involved in nervous system processes (116 genes), particularly those that act through G protein-coupled receptor activity (140 genes) and neurotransmitter receptor activity (36 genes). This heightened allocation to signaling *between* cells is in contrast to the downregulation we saw next of processes *within* cells.

We found the downregulated archetype involves growth processes that run the day-to-day metabolic and protein turnover processes inside the cell. Those having reduced expression include cullin-5, a ubiquitin protein ligase, and atp-2, the beta subunit of ATP Synthase. More broadly, they include the pathways for mTOR signaling, mRNA surveillance, and protein degradation (proteasome) that regulate growth, as well as glycolysis/gluconeogenesis, the TCA cycle, and oxidative phosphorylation that power this growth. Also downregulated are components of ATP Synthase, which is notable given its importance both in aging mice [26] and in the regulation of mTOR signaling and transcriptional drift [27].

Overall, *C. elegans*’s large-scale transcriptional drift appears to take a quasi-beneficial or adaptive path where synthesis of high-biomass pathways consisting of long-lived proteins are made early in life (thus downregulated with age), while signaling processes needed to sense the environment and coordinate beneficial actions are relatively overexpressed later in life.

## Transcriptional drift of individual genes

So far, our analysis is coarse-grained, covering the average behavior of the two archetype profiles. Here, we now focus on some of the individual genes that make up each archetype. Rangaraju et al. reported their data on transcriptional drift for each particular gene as TD_ratio_ = cpm_*x*_/cpm_0_ where cpm (counts per million reads mapped) refers to the expression level of a gene as measured through RNA-seq, and where the subscripts indicate day number x starting from day 0, which is the first day sexual maturity is reached [14]. Our purposes here are best served by normalizing relative to day 0, cpm_*x*_ − cpm_0_, and relative to the mean, to prevent overemphasis on outliers, so instead we use

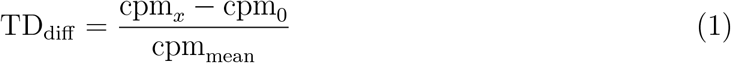

where cpm_*mean*_ is the mean cpm across the time points we analyzed.

## A proposed mechanism: The Cumulative Factor Model

What mechanism might explain such largescale coordination of up-regulated and downregulated protein levels with age? Here, we propose a minimalist model in which simply the concentration [*f*] of a single underlying molecular factor *f* – say, some ligand or protein – rises passively with age. This could result from some age-related decline, such as in proteostasis [28] or metabolism [2], or it could represent an age-related program that changes proteome composition and energy expenditure in a way that improves fitness [29, 30]. Regardless of the exact mechanism, a single such factor would be sufficient to drive concerted expression levels of large subsets of the genome [31].

Here’s how it could work. Suppose the factor concentration [*f*] accumulates linearly with age. If *t* is the cell’s adult age (the time since completion of larval development), then the concentration of *f* at time *t* is:

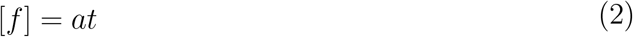

where *a* is a constant rate of accumulation of the factor.

Next, we describe how the cell turns that aging signal into a modulation of gene expression levels in the model. We suppose that sensory genes are precursors to genes that respond to stress [32]. The cell uses these sensory genes to detect the type of stress (start/end of starvation, oxidative, osmotic, DNA damaging stress, etc.), and signaling initiates cellular responses to counteract the stress as an attempt to reestablish homeostasis [33]. Thus, we make the assumption that signal genes correlate with stress genes.

In a young cell, we expect that the number of mRNAs and proteins involved in growth are in some optimal balance relative to those involved in signaling/stress. The data indicates that at time *t* = 0, in young cells, we have an initial level of signal genes *s*_0_, averaged over all the corresponding proteins in that class. We suppose that *s*_0_ *< g*_0_ since a young cell has seen little stress yet and is poised to grow. However, cells will naturally experience stress throughout their day-to-day activities. This creates gradual change in transcriptome regulation and proteotoxic stress which the cell must adapt to, resulting in a necessary increase in signal-related gene expression [14, 32].

The cell can detect its age by monitoring [*f*] through Langmuir-type binding of *f* to a stress-sensor biomolecule. Thus, the number of signal mRNAs, *s*, will be

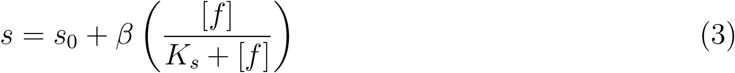

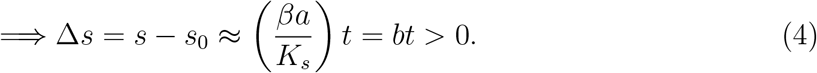

where *s*_0_ is the initial concentration of the average signal mRNAs, *K*_*s*_ is a binding association constant, *β* gives the number of molecules binding and *b* = *βa/K*_*s*_ is the slope of the time dependence in units of *s* per unit time. Mathematically, *s*_0_ is the minimum value of *s*, at time zero and the signal gene expression is an increasing function of age. Because these time courses are observed to be linear, we are able to make the approximation that *a « K*_*s*_. So we can fit the experimental data with a single parameter *b*, which gives mechanistic insight because it is proportional to the average number of mRNA copies made for the signaling/stress sub-genome. Note that Δ*s* is a measure of *TD*_diff_ for signal genes.

The same mechanism applies to the growth genes, where *TD*_diff_ is described by Δ*g*:

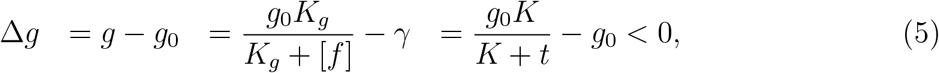

Here, *g*_0_ is the maximum value of *g*, which occurs at the youngest age since *g* is a decreasing function of age. *K* = *K*_*g*_*/a* is the binding association constant for growth genes. Since the levels of the growth genes are not linear in age (unlike the stress genes), we now require two parameters, *g*_0_ and *K*, to fit the experimental data. For fitting the experimental curves, we first rescaled the cpm at each time point of each gene by its mean expression across time. This allowed for better comparisons between genes without changing the overall shape of the data.

### Fitting to Experimental Data

We now use the experimental data to assess these linear and Michaelis-Menten binding mechanisms. We plotted best-fit curves to the average value of the change in cpm of sensory genes using equation 4. This allowed us to approximate a value for [*f*] since it is directly correlated with *b*, the resulting best-fit coefficient. This was then used to fit equation 5 to the average value of the change in the cpm of all growth genes using equation 5. As demonstrated in figure 3, both equations give food fits for the observed cpm with *R*^2^ values of *>* 0.99. Taken together with the observation that drugs and longevity genes can broadly delay this drift [14], this supports the hypothesis that signal and growth genes are coordinated – rising and falling together under shared control – because each group’s behavior is characterized by their dependence on a shared variable, [*f*] = *at*.

**Figure 3.**
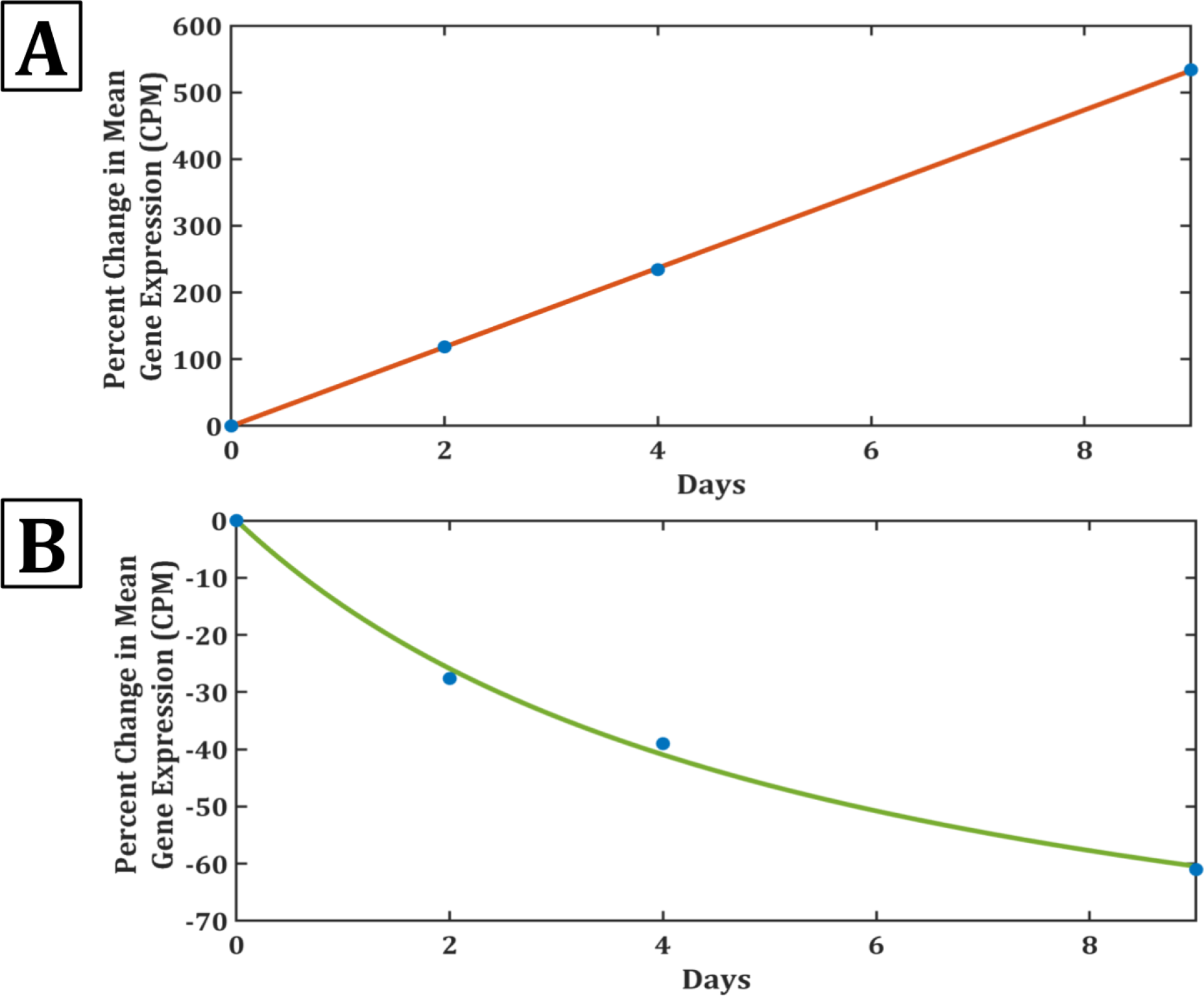
The Cumulative Factor model captures the linear increase in upregulated expressions and the Michaelis-Menten decrease in downregulated expressions. Best-fit regressions to the data using the equations in the text. The average value of the percentage change in cpm across all genes in each subset is plotted in blue. Plot A models the sensory genes with an *R*^2^ of 0.9999 and plot B models the growth genes with an *R*^2^ of 0.9964.

**Table 2:**
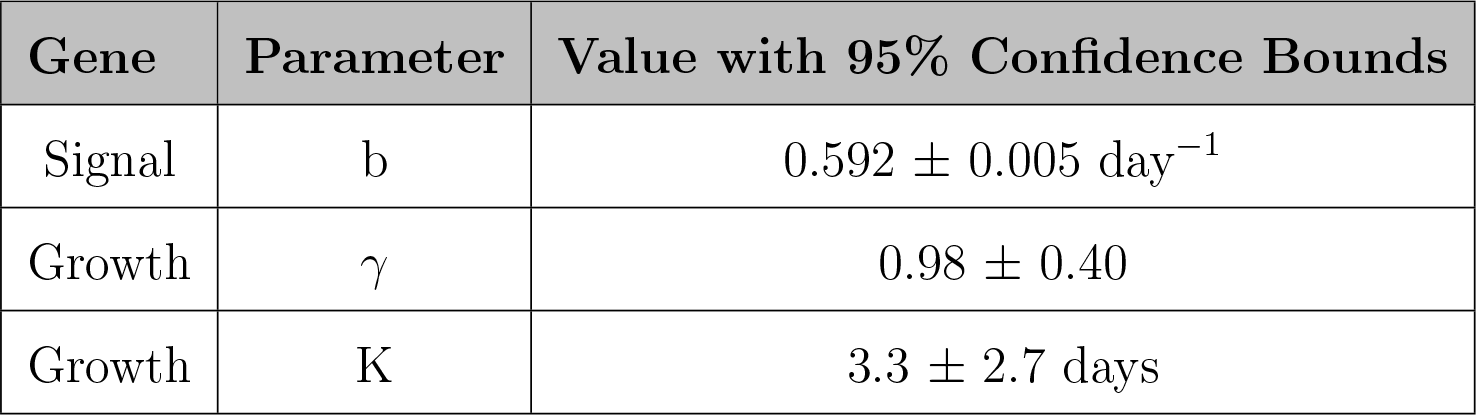
Parameter values for equation fits.

| Gene | Parameter | Value with 95% Confidence Bounds |
| --- | --- | --- |
| Signal | b | $0.592 \pm 0.005 \text{ day}^{-1}$ |
| Growth | $\gamma$ | $0.98 \pm 0.40$ |
| Growth | K | $3.3 \pm 2.7 \text{ days}$ |

## Discussion

We have distilled the complexity of *C. elegans* aging to an elementary form, showing how a single factor [*f*] accumulating linearly with age is capable of regulating the concerted drift observed in a large fraction of the aging transcriptome. Both upward and downward expression follow Michaelis-Menten or Langmuir-like binding forms, with the activating binding-action of the sensor archetype being less saturated - and thus more linear - than the saturating downward growth archetype. While the physical identity of this regulating factor [*f*] remains unknown, there are a few key possibilities.

Firstly, age-related drift in gene expression has been associated with changes in the levels of master regulators such as daf-16 and skn-1[32, 34] – known to control growth and stress resistance [29] – as well as to changes of regulatory miRNAs that impact mRNA turnover [11]. While the activity of these regulators [34] may individually not be as smooth as the genome-wide patterns seen here [14], they may collectively shape – and be responsive to – the underlying changes captured by our factor [*f*].

Secondly, our factor [*f*] could reflect the accumulation of a more distributed, bottom-up loss of information. For example, something as basic as making mRNA molecules and their protein products in the correct ratios to form a functional multi-protein complex or pathway fails with age [15, 35, 27, 36, 37]. This loss of coordination could arise from the accumulation of random changes in the epigenome that impact mRNA production (mRNA-first stoichiometry loss) [14] or may result from less efficient or spatially-localized translation and assembly of protein complexes (protein-first stoichiometry loss) [37]. Any protein subunits made in excess of the functional ratio would need to be stabilized and degraded, creating a proteostasis burden that scales with growth rate. Adaptation to this loss of biological coordination would favor the rise of sensing/stress genes and the decline in growth genes seen here.

Lastly, it should be acknowledged that individual cell types undergo unique aging trajectories at the gene and pathway levels [32]. Each cell appears to be adapting to stresses unique to their cell type, activating different sets of stress response genes that delay their aging decline. For example, neurons upregulate protective skn-1 target genes. At the same time, they strongly downregulate respiratory metabolism [32], an action that may amplify cell-wide transcriptional changes [38]. In contrast, the rise of heat shock proteins is shared across cells, suggesting that protein folding and assembly is a fundamental stress closely related to our factor [*f*] [32]. For deeper insights into the mechanism and nature of this factor, we advocate for experiments applying external variations, such as temperature, osmolarity, and pH that cause changes in the transcriptome [39, 9].

## Acknowledgments

This work was supported by the Laufer Center of Physical and Quantitative Biology at Stony Brook University and by the National Institute of General Medical Sciences MIRA Program (R35 GM122561). We thank Dr. Michael Petrascheck for his valuable insights and assistance with interpreting the data from Rangaraju et al.

## SI

Archetype Analysis

NMF is well-known in the mathematical literature. We have recently develop a variant of NMF relevant for clustering gene expression trade-offs as a function of external variables [21]. Given an *N × M* data matrix *V*, NMF generates two new lower-rank matrices, *W* and *H* such that:

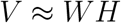

where *W* is an *N × k* matrix representing coefficients of each signal contribution to the *k* archetypes, and *H* is a *k × M* matrix representing the weights of each archetype needed to approximate each sample. As the relative *H* weight of a given archetype goes up, necessarily the total relative weights of the others go down. For a visual representation of the relationship between NMF and multi-signal trade-offs see Figure 1 of [21].

To reduce noise, we applied this methodology to the observed trial-averaged gene expression values. To remove bias towards highly-expressed genes, we then re-scaled these values by the mean expression (over time) of each gene. We then filtered for genes with remaining variance between 0.1 and 0.2 to remove additional biases towards outlier genes and to limit the RAM utilization of archetype analysis. As a final pre-processing step for archetype analysis, the expression levels of each gene were normalized so that the total expression at each time point equaled 1 [40, 21]. Using the eigenvalue elbow criterion on the resulting data matrix, we determined that two signatures (the minimal number possible) sufficiently represented the *C. elegans* transcriptome. These archetypes represent collections of genes that show either monotonic increases or decreases in abundance with age, measured on days 1, 3, 5, and 10 of adulthood.

**Figure A1.**
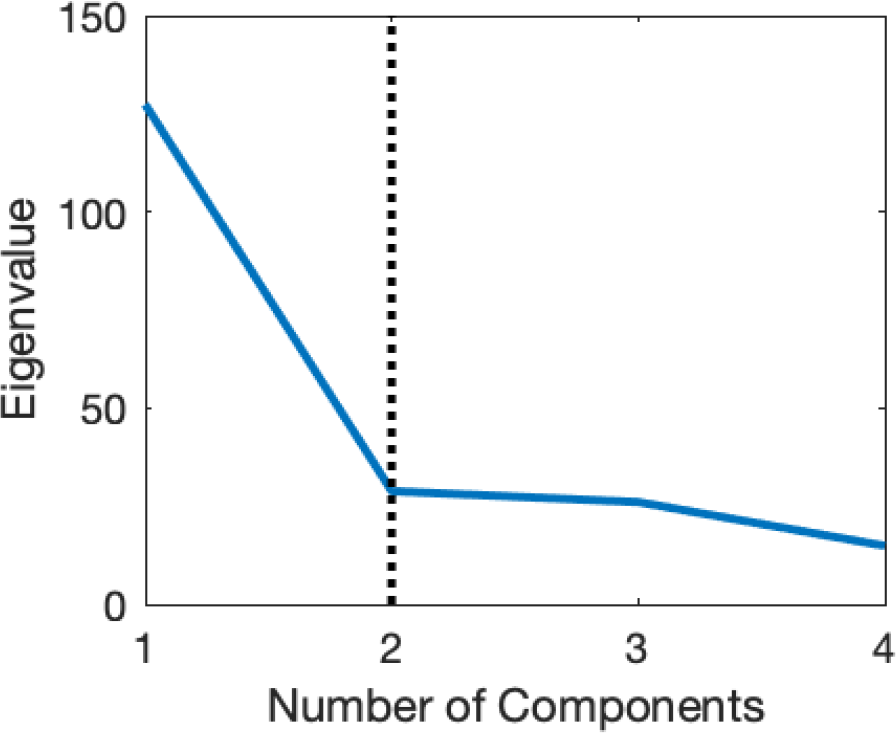
Eigenvalues of the *C. elegans* transcriptomic data. The dotted line reflects the “elbow” of the eigenvalue spectrum, suggesting that 2 components should be retained for downstream analysis.

